# Revealing sphingolipids composition in extracellular vesicles and paternal β-cells after persistent hyperglycemia

**DOI:** 10.1101/2024.01.06.574464

**Authors:** Magdalena E. Skalska, Martyna Durak-Kozica, Ewa Ł. Stępień

## Abstract

Extended periods of hyperglycemia (HG) can lead to metabolic disorders of sphingolipids (SPs) and their subsequent accumulation in cells. This accumulation can trigger a range of complications, including kidney and neurodegenerative diseases. In our study, we compared the levels of selected ceramides (CER), hexosylceramides (HexCER), and glycosphingolipids (GSLs) in potential HG biomarkers - extracellular vesicles (EVs). These EVs were derived in vitro from human β-cells cultured under both normoglycemic and high-glucose conditions (HG). We utilized Time of Flight – Secondary Ion Mass Spectrometry (ToF-SIMS) for SPs analysis. Our results confirmed that the lipid profiles of these three groups differ between large and small EVs, with some SP lipids being more enriched in EVs compared to cells. Interestingly, our study revealed that HG only regulates the lipid content from the glycosphingolipid group in relation to normoglycemia. Collectively, our findings underscore the potential applications of ToF-SIMS in characterizing the impact of different culture conditions on lipid levels. As far as we know, our study is the first which employs ToF-SIMS in analyzing the effects of HG on SP levels in EVs and their parental β-cells.

**Graphical abstract:** 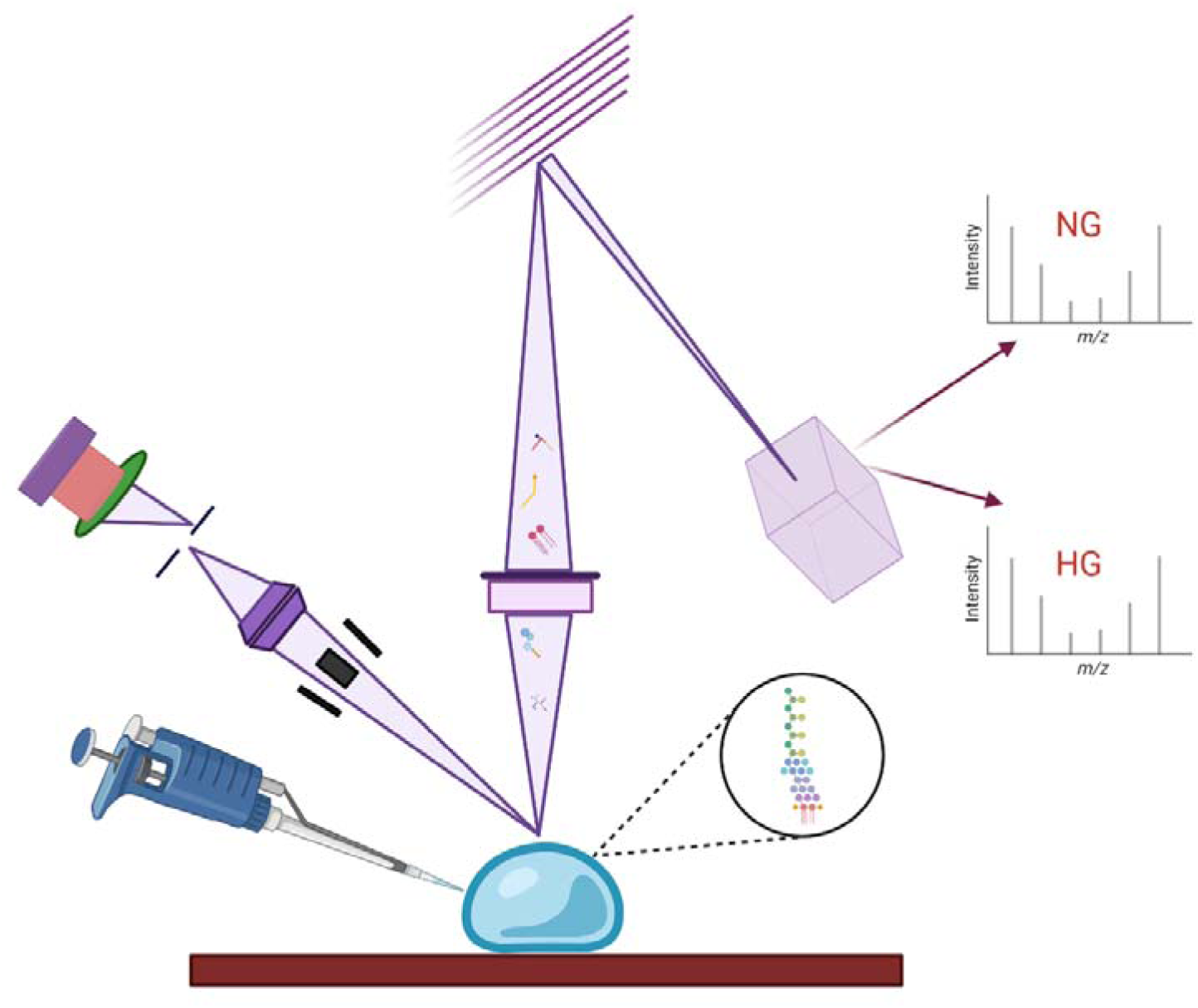

**Highlights:**

- ToF-SIMS effectively compares glycosphingolipid content in cells and EVs.
- Analysis reveals lipid profile differences between EVs and their parent cells.
- Hyperglycemia alters glycosphingolipid content in cells and EVs per ToF-SIMS.

## 1. Introduction

Sphingolipids (SPs), including ceramides (CERs), hexosylceramides (HexCERs) and glycosphingolipids (GSLs), represent a ubiquitous and structurally miscellaneous class of lipids involved in physiological and pathological processes, possessing both signaling and structural properties [1,2]. Lipid membrane composition is varied in different cell compartments, and has impact on such cell membrane properties, charge and protein corona [3]. Hyperglycemia (HG) is an example of a physiological condition initiated by elevated blood glucose levels, which, when out of control of the sensitive fuse that is the hormone insulin, causes metabolic stress, which, if it continues chronically, can cause pathological conditions. In fact, in most patients having glucose intolerance, HG goes silent early on and is often not detected until complications develop. What is known, however, is that the first stage in the development of complications is elevated glucose level for a dangerous period, followed by insulin resistance, eventually leading to type 2 diabetes mellitus (T2DM) [4]. An important role in glycemic control and HG prevention is played by pancreatic β-cells, which can compensate for high glucose concentrations by increasing insulin secretion [5]. This condition, in which HG is suppressed by insulin overproduction, can persist for years and may never develop into overt diabetes [6,7]. And yet, over time, the ability of β-cells to produce high levels of insulin may be diminished due to both abnormal cell signalling and increased apoptosis [8]. In this case, HG modifies the molecular composition of pancreatic β-cells that determine metabolic processes, accelerating the course of β-cells to failure and disease progression [9]. Indeed, SPs have been shown to actively participate in the HG-induced process, mediating the loss of insulin sensitivity, promoting the characteristic diabetic pro-inflammatory state, and inducing cell death and dysfunction of important organs such as the pancreas and heart [10,11]. However, given the high complexity of SP metabolism, it follows that its regulation may also be very complex or multifactorial.

Time of Flight – Secondary Ion Mass Spectrometry is an example of an advanced physical method that can be successfully used in metabolomic and lipidomic researches. This technique has been successfully used for the characterization of lipids in biological samples and comparative analyzes of cells, tissues, organs, and even organisms [12–14]. In this study, we would like to focus on SPs composition in β-cells and their residues and metabolites secreted in the form of extracellular vesicles (EVs) using ToF-SIMS.

EVs are spherical nanostructures surrounded by a lipid bilayer ranging in size from 30 to 1000 nm. EVs can be divided into several subpopulations that differ in biogenesis, size, cargo and function [15]. Currently, the main EV populations recognize large EVs (150 – 1000 nm) and small EVs (50 – 150 nm) [16]. Their size is of great importance because it can be used to assume the biogenesis and function of EVs, while large EVs are directly secreted from the cell membrane, small EVs accumulate in the lumen of late endosomal compartments and are released into the extracellular space after endosome fusion with the cell membrane [17,18].

It is believed that the most important function of EVs is the transfer of intercellular information in the form of nucleic acids (miRNA), proteins and lipids. They contain characteristic cell-derived molecules that reflect the intercellular status of their parent cell. Thanks to this cargo, they affect the physiological and pathological reactions of cells receiving the transmitted data, thus taking an active part in maintaining the homeostasis of the biological system by transmitting information, but also expelling unnecessary material [19]. Research results of the last decade have revealed that EVs are released from various cells, and their cargo is involved in various disorders, mainly cancer, neurological and metabolic, e.g. diabetes [20]. In biomarker studies but also to understand EV signalling properties, the molecular characterization of EVs obtained from different cellular models (cell cultures) is essential in both basic and preclinical research [16,21,22]

The lipid composition of EVs is reflective of the cells from which they originate [23]. Consequently, it can be inferred that the composition of sphingolipids (SPs) in EVs would vary based on the cell culture they are derived from, as well as the conditions under which they were cultured. Past studies have indicated that the molecular composition of both cells and EVs can be altered by external culture conditions [24]. However, it remains to be seen whether an elevated glucose concentration in the external environment can modify the profile of ceramides (CERs), hexosylceramides (HexCERs), and glycosphingolipids (GSLs) in the composition of β-cell membranes and EVs.

In this study, we employ ToF-SIMS mass spectrometry to compare the profiles of selected sphingolipids (CER, HexCER and GSL) in EVs samples derived from a β-cell line with the compositions cells cultured under HG conditions.

## 2. Materials and methods

### 2.1. Cell cultures

The insulin-releasing pancreatic β-cell line (1.1B4, Merck KGaA, Sigma Aldrich, Darmstadt, Germany) was cultured in a 175 cm^2^ cell culture bottles and maintained in RPMI 1640 medium (cat. No. 11879020 Gibco for normoglycemia and hyperglycemia, 21875158 Gibco for medium hyperglycemia), supplemented with 10% Fetal Bovine Serum (FBS; cat. No. 16000036, Thermo Fisher Scientific, Waltham, MA, USA), 2 mM L-glutamine (cat. No. G7513 Sigma-Aldrich, Darmstadt, Germany) and antibiotics - 100 µg/ml streptomycin with 100 units/ml penicillin (cat. No. 15140122 from Sigma-Aldrich, Darmstadt, Germany). The cells were grown as monolayers in an incubator under conditions of a humidified 5% CO_2_ atmosphere and 37 °C until they reached about 80% confluence. The culture was maintained by changing the medium and passaged with trypsin (Trypsin-EDTA solution, cat. No. 25200056 Thermo Fisher Scientific, Waltham, MA, USA). The experiment involved growing cells in three different glucose concentrations: normoglycemic control (NG) with 5 mM D-glucose (cat. No. G8270 Sigma-Aldrich, Darmstadt, Germany), medium hyperglycemia conditions (MDM) with 11 mM D-glucose, and hyperglycemia (HG) with the highest concentration of glucose at 25 mM D-glucose. After three passages, the cells were starved for 24 hours by replacing the medium with a serum-free one. Once the fasting period was over, the culture bottles were mechanically shaken, and the medium was collected. The collected fluid was then subjected to subsequent centrifugation and low-pressure filtration dialysis to concentrate and isolate EVs. The medium was collected from 12 culture bottles for all culture conditions. From each 4 cell culture bottles, the medium was collected in a separate container (approximately 120 ml), yielding three biological replicates for NG, MDM and HG conditions.

### 2.2. Cells Harvesting

To compare the molecular composition of lipids, cells were collected through trypsinization after a serum starvation period in NG, MDM, and HG. These cells were then suspended in PBS and frozen at −80 °C. This was done to enable simultaneous analysis with isolated EVs.

### 2.3. EVs Isolation

The collected volumes of β-cell conditioned medium were prepared by two centrifugations to remove intact cells, cell debris, and apoptotic bodies (**Fig. 1**). The cell media samples were successively centrifuged at 400 x *g* (10 min) and 3,000 x *g* (for 25 min), both at 4 °C. After that, each 120 ml sample was filtered under Low-Pressure Filtration Dialysis (L-PFD) using a cellulose dialysis membrane (cat. No. 131486, Spectra/Por Biotech) with a molecular weight cut-off (MWCO) of 1000 kDa (width: 16 mm, diameter 10 mm). To accelerate the filtration process, a reduced pressure of −0.2 bar was applied in the system, which helped in obtaining the final sample volume of about 1.5 ml [25,26]. Finally, the membrane was then washed with 10 ml of deionized water.

**Fig. 1.**
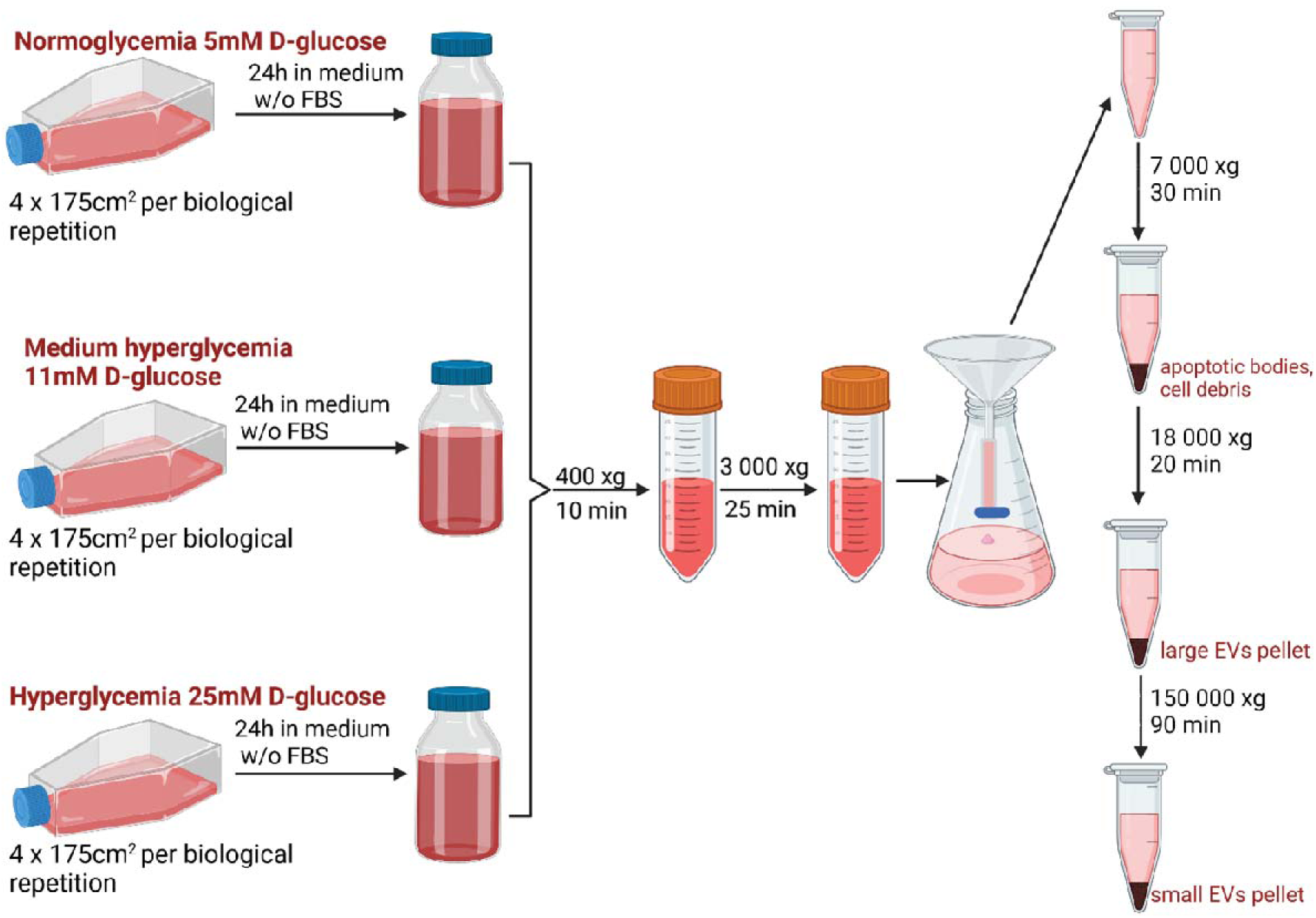
The scheme for the extracellular vesicle isolation procedure.

The EV suspension was treated with a differential centrifugation method. In the first step, the samples were centrifuged at 7,000 x *g* for 30 minutes at 4 °C and then at 18,000 x *g* for 20 minutes to isolate the **large EVs**. Next, the resulting supernatant was transferred to 1.5 ml top-opening centrifuge tubes and centrifuged for 1.5 hours at 150,000 x *g* at 4 °C (Sorvall MX 150+ Micro-Ultrawiring, Thermo Fisher Scientific, Waltham, MA, USA) to obtain the subpopulation of **small EVs**.

Pellets were resuspended in 50 µl PBS (cat. No. 10010023 Thermo Fisher Scientific, Waltham, MA, USA) in triplicate and stored at −80 °C for further characterization and ToF-SIMS analysis.

### 2.4. Antibodies for flow cytometry measurement

The small and large EVs samples were characterized using FITC Annexin V (BioLegend, cat. 640906), antibodies Alexa Fluor 700 anti-human CD63 (BioLegend, cat. 353024) and PE anti-human CD9 (BioLegend, cat. 312106).

### 2.5. Spectral Flow Cytometry

Both small and large EVs were isolated from 120 mL of conditioned media per sample, which were then divided into 4 groups. The EVs were then incubated with Annexin V and antibodies against CD63 and CD9 for 45 minutes on ice. The EVs’ samples were analyzed using the ID7000 apparatus (spectral flow cytometer, SONY), and the sample volume was set to 100 μL. Before to analysis, the Sony ID7000 was calibrated using alignment checks (Sony Biotechnology Inc AlignCheck Flow Cytometer Alignment Beads 10^7^/mL 10 µm, 2 mL, cat. no AE700510) and the 8[peak performance beads (Sony Biotechnology Inc 8 Peak Bead cat. no AE700522, 10^7^/mL 3.1 µm, 5 mL cat. no AE700510), following the instrument supplier’s guidelines.

### 2.6. Transmission Electron Cryomicroscopy (cryo-TEM)

Approximately 3□μL of the sample solution were put on newly glow-discharged TEM grids (Quantifoil R2/1, Cu, mesh 200) and plunge-frozen in liquid ethane using the Vitrobot Mark IV (Thermo Fisher Scientific). The following parameters were used: humidity of 95%, temperature of 4□°C, and blot time of 2□s. The frozen grids were stored in liquid nitrogen until they were cut and loaded into the microscope. Cryo-EM data were collected at the National Cryo-EM Centre SOLARIS (Kraków, Poland). Movies (40 frames each) were captured using the Glacios microscope (Thermo Fisher Scientific) with a 200□kV accelerating voltage, 190 kx magnification and corresponding to a pixel size of 0.74□Å/px. The direct electron detector Falcon4 was used to obtain data, and it was operated in counting mode. The images were exposed to a total dose of 40 e-/Å2 total dose each (measured *’on vacuum’*). Images were acquired at under-focus optical conditions.

### 2.7. Transmission Electron Microscopy (TEM)

To perform negative staining, a copper grid coated with formvar and carbon was used. The samples were incubated on the grids for 3 minutes, after which the excess was removed with a piece of filter paper. The grids were then washed with two drops of distilled water and uranyl acetate. After rinsing, the grids were left on a drop of uranyl acetate for one minute and then dried in the air. For observation, the JEOL JEM 2100HT electron microscope (Jeol Ltd, Tokyo, Japan) was used at an accelerating voltage of 80 kV. Images were taken using a 4kx4k camera (TVIPS) equipped with EMMENU software *ver*. 4.0.9.87. The image analysis was performed using the Fiji program. The histogram was made based on 100 EVs counts.

### 2.8. ToF-SIMS

Time of Flight - Secondary Ion Mass Spectrometry (ToF-SIMS) is a surface analytical technique based on the emission of secondary ions from the examined surface due to a surface bombardment with heavy and high-energy primary ions. In the case of biological samples, ionized clusters of bismuth or fullerene (Bi_3_^+^, C_60_^+^) are typically used as they minimize damage to the structure being tested while allowing the detection of larger molecules. The spectrometer employs a time-of-flight analyzer with high mass resolution (up to *m/*Δ*m* = 10,000) and high transmission. It measures the time of flight of emitted secondary ions through a vacuum tube, including ionized whole biomolecules, ionized fragments of these biomolecules, or newly formed ionized conjugates. Some of these ionized molecules are characteristic fragments of biomolecules that are components of tested biological structure, such as amino acids, lipids or metabolites, which enable the qualitative characterization of the sample surface [27]. It is possible to carry out the described analysis in so-called *static mode*. This mode uses a dose density threshold of primary ions deposited on the tested surface to ensure that the surface is not damaged and the components are not significantly delocalized and fragmented in the sample volume [28].

A significant advantage of the ToF-SIMS technique for biological samples is its ability to perform comparative analyses of experimental samples with control samples of known physical and chemical properties. This is done by assessing the intensity of peaks characteristic of the biomolecules of interest, eliminating the need to extract these compounds from the biological structure or to label these structures in the sample. This feature is particularly important for lipids that are susceptible to oxidative processes. However, the requirement of high vacuum measurement in an analytical chamber poses a challenge for biological research, adding a layer of complexity to the procedure of preparing biological samples.

### 2.9. ToF-SIMS measurements

For ToF-SIMS measurements we used silicon wafers (cat. No. 647780, Sigma Aldrich, St. Louis, MO, USA) with dimensions of 1 x 1 cm^2^ [29]. Before depositing EVs and cells, each substrate was sonicated in toluene (cat. No. 244511, Sigma Aldrich) and absolute ethanol (99%, cat. No. 396480111, POCH) for 10 min in an ultrasonic bath. The substrate was then rinsed with plenty of deionized water and dried with N_2_. We used a ToF-SIMS 5 instrument (ION-ToF GmbH, Münster, Germany) with a Bi_3_^+^ liquid metal ion gun (30 keV) as the primary ion source to record data in the mass range up to 900 Da. The experiments were performed in the static spectrometer mode with a current equal to 1.10 pA, and a low-energy electron gun was used to neutralize the charge formed on the sample surface. All mass spectra were measured simultaneously and under the same conditions using a single sample holder.

**Fig. 2** presents the workflow of the process of preparing data from spectra for comparative analysis. Three types of samples were prepared from each culture condition: cells, large EVs, and small EVs in three biological replicates, resulting in 27 samples for ToF-SIMS analysis. For each sample, a volume of 30 µL deposited on the silicon surface was prepared.

**Fig. 2.**
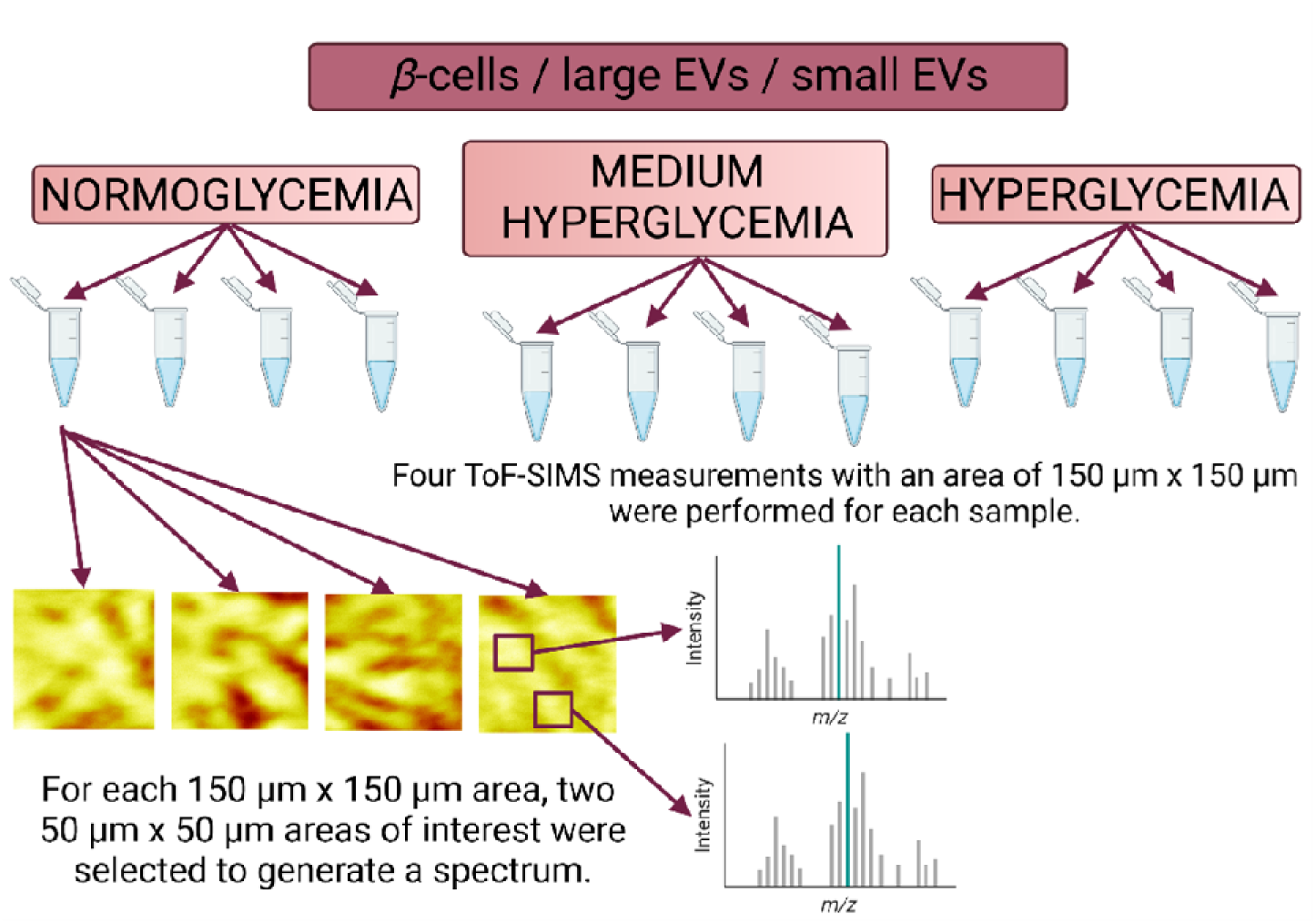
Workflow of obtaining spectra for comparative analysis.

Three surface measurements were taken for each sample, with a size of 150 × 150 µm^2^ for positive ions (125 x 125 pixels). For each measurement, two areas of 50 x 50 µm^2^ were selected, referred to as *Region Of Interest* (ROI). Before this, 2D emission maps were examined for characteristic ions for phosphocholine fragments (*m/z* 184.2), which helped to eliminate local artefacts.

The normalization of raw ToF-SIMS data is typically performed to mitigate artefacts induced by topographic, apparatus, or matrix effects [30–32]. To obtain semi-quantitative information from the measured data, normalization was done using the total dose deposited on the examined surface, and each spectrum was calibrated using signals for positive ions: *H*^+^, *H*_2_^+^, *CH*^+^, *CH*_2_^+^, *CH*_3_^+^ and *C*_3_*H* ^+^. The obtained spectra were further analyzed, which included the identification of characteristic lipid peaks such as protonated ions, adducts, and pseudo-molecular ions. It should be noted that different molecular formulas may correspond to the same molecular weight. Only peaks with a high likelihood of matching the structures of interest and those that have been previously described and used in the ToF-SIMS analyses were included in the comparative analysis (**Supplementary A**). The data was analyzed using SurfaceLab software (*ver.* 7.2), which enabled re-editing of the measurements taken. Additionally, the academic version of OriginPro (*ver.* 9.8.0.200, 2020b) was used for further analysis.

## 3. Results

### 3.1. Spectral Flow Cytometry

We conducted a flow cytometry analysis on our isolated samples to determine the different types of EVs present. To identify ectosomes, we used Annexin-V as a marker and found a positive signal in both small and large EV samples (**Fig. 3, 4**). For small EVs and small ectosomes, we used CD9 and CD63 as markers and detected positive signals in both samples (as shown in **Fig. 3, 4**). Interestingly, large EVs showed higher expression of CD63 and CD9, while annexin V was more prevalent in the population of small EVs. As demonstrated in **Supplementary B**, pure PBS yielded only a minimal signal.

**Fig. 3.**
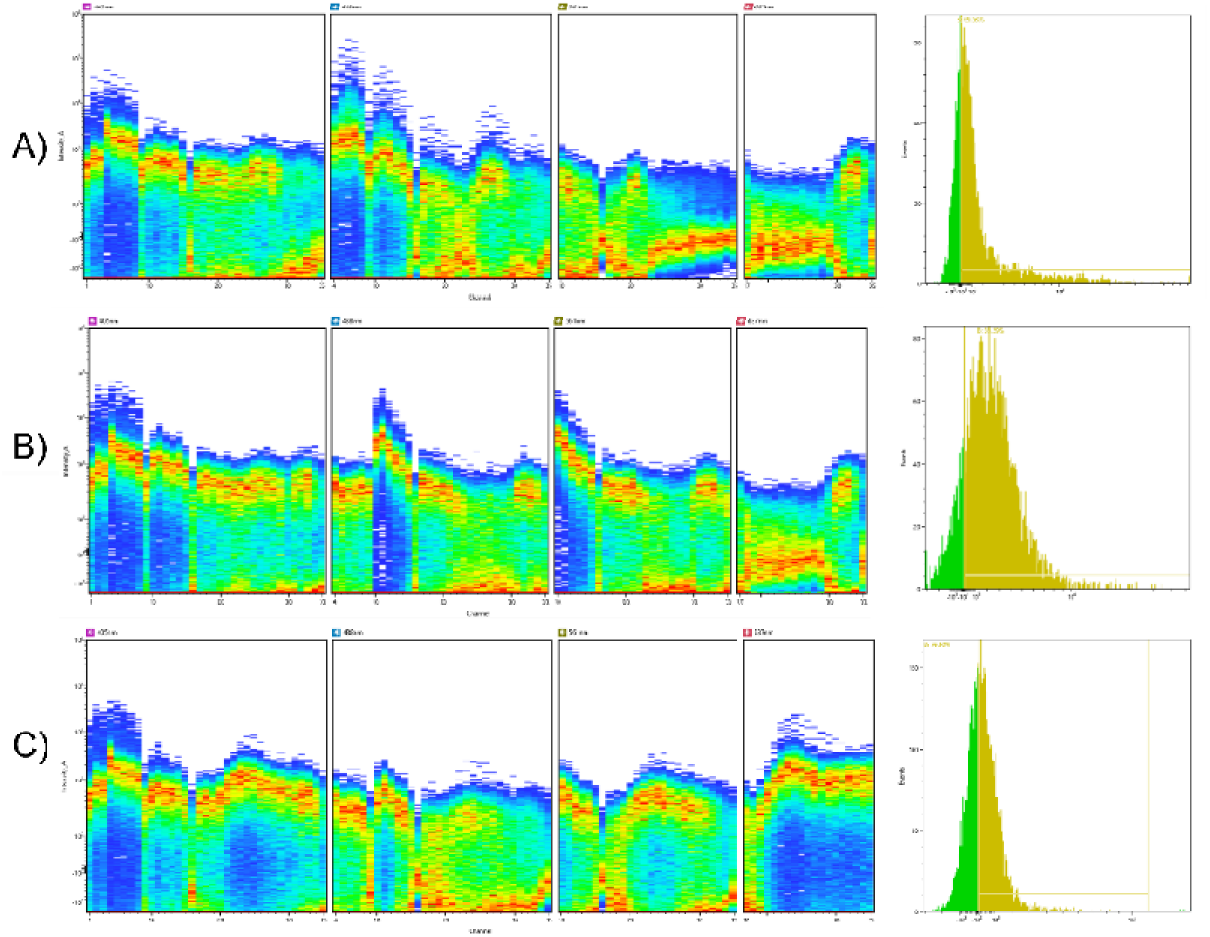
Ribbon plots for stained large EVs with histograms: A. FITC annexin V, B. PE CD9, and C. AF700 CD63.

**Fig. 4.**
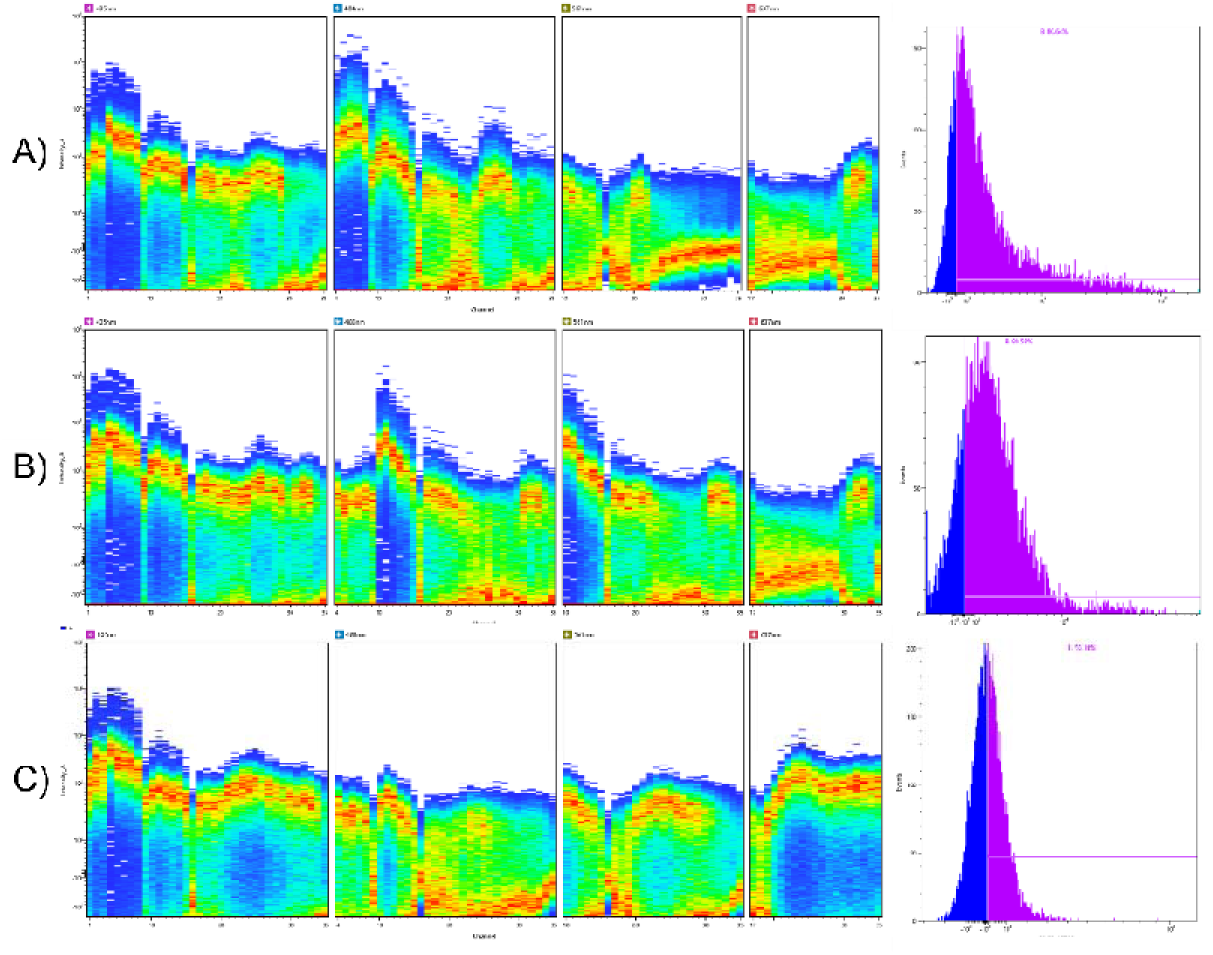
Ribbon plots for stained small EVs with histograms: A. FITC annexin V, B. PE CD9, and C. AF700 CD63.

### 3.2. TEM and cryo-TEM

In **Fig. 5 A, B)**, an empty copper grid is demonstrated, which confirms the successful isolation of EVs. The average size of large EVs is around 150 nm, as illustrated in **Fig. 5 C** and **5 E**, while small EVs have a mean size of approximately 60 nm, as displayed in **Fig. 5 D** and **5 F**. These findings not only confirm the distinct sizes and shapes of both EV populations but also provide valuable insights into their relative abundance, which suggests a higher concentration of small EVs.

**Fig. 5.**
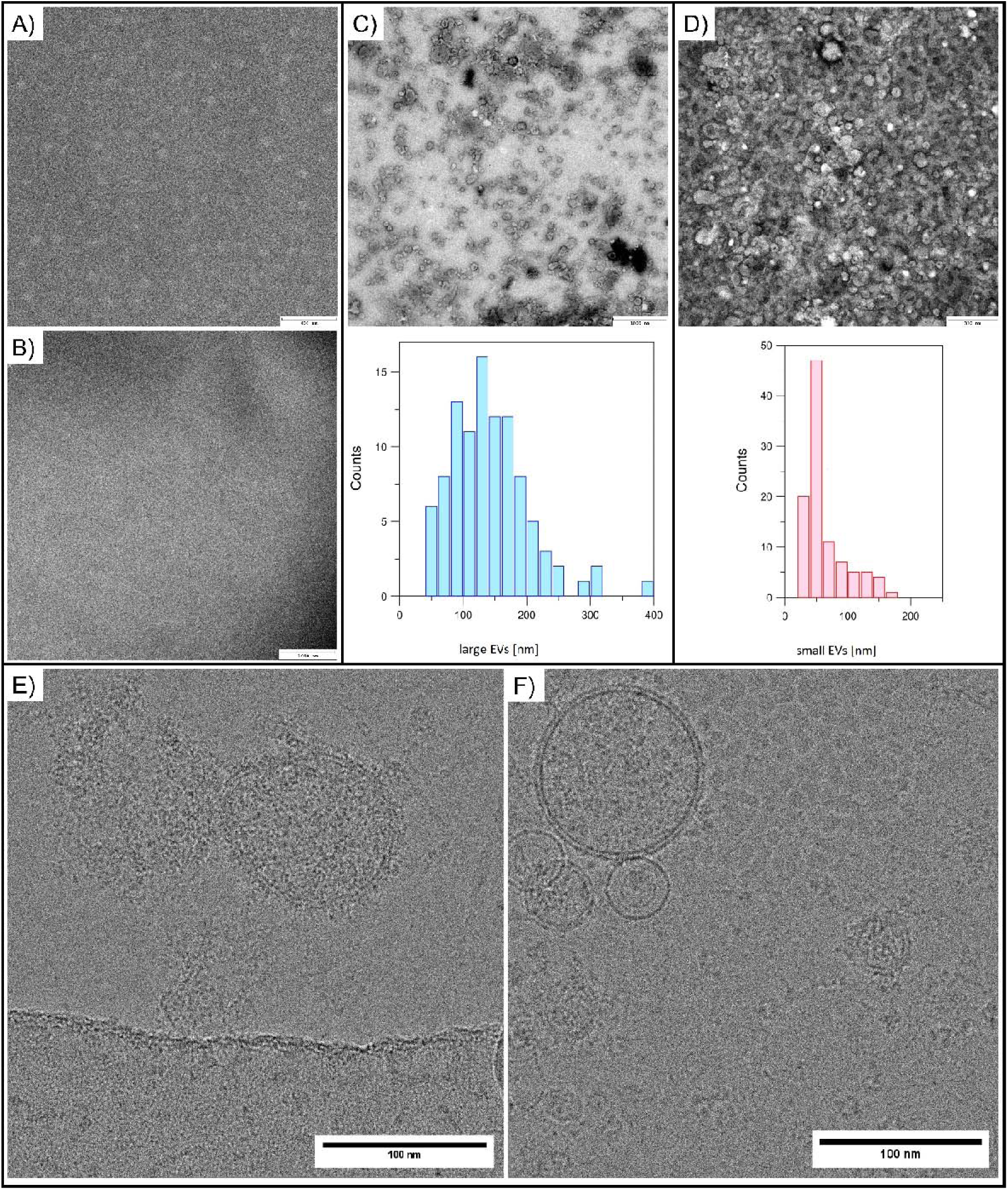
A) The copper grid before contrasting, B) the copper grid after contrasting, C) the image and size distribution histogram for the population of large EVs, D) the image and size distribution histogram for the population of small EVs, E) the cryo-TEM image of large EVs, F) the cryo-TEM image of small EVs

Images taken using cryo-TEM microscopy (**Fig. 5 E, F**) reveal the double membrane encapsulating the EVs, as well as the cargo inside the vesicle. Specifically, in the case of larger EVs (as seen in **Fig. 5 E**), a crown-like structure can be observed on their surface. Further studies using TEM and cryo-TEM have verified that the EV samples are free from any cells, cellular organelles, or other cellular debris.

### 3.3. ToF-SIMS analysis

The molecular composition of EVs is thought to reflect the composition of secretory cells and is influenced by various factors [33]. EVs transfer information to other biostructures due to their structure and the contained cargo [34].This information is encoded in the presence and quantity of building blocks, including approximately 9,769 proteins, 1,116 lipids, 3,408 mRNAs and 2,838 miRNA [35]. In the present analysis, three groups of lipids were examined: ceramides (CERs), hexosylceramides (HexCERs), and glycosphingolipids (GSLs).

These lipids are part of the cellular endosomal complex that orchestrates and directs the endogenesis pathway within the cellular system. They play a crucial role in influencing the formation of membrane-like structures secreted by cells, mainly CERs and other SPs [36]. They also impact the content and destination of the endosome [37–39]. C18:0-Cer and C24:1-Cer molecules are the dominant CERs present in small EVs, which facilitate the membrane folding and differentiation of secreted small EVs [40]. GSLs exist as lipid rafts that can envelop and penetrate the surface membrane or manifest as a flat lipid rafts. These structures are enriched with specific types of lipids, such as sterols, SPs, and glycosylphosphatidylinositols. Through structural changes in lipid raft components and protein sorting and the formation of complexes with tetraspanins or flotillins, they play a pivotal role in endosomal biogenesis [41,42].

Previous studies have suggested that SPs may play a role in the development and progression of various pathologies, including obesity and type 2 diabetes mellitus (T2DM) [43]. Therefore, recognizing positive or negative correlations between SP levels and outcome conditions may contribute to identifying useful and early indicators for these pathologies.

The ToF-SIMS technique was used to investigate the normalized average intensity of characteristic peaks changes in the case of β-cells, large and small EVs secreted by these cells.

#### a) Ceramides (CERs)

***CERs*** are lipids composed of a long-chain or sphingoid base linked to fatty acids by amide bonds. Free CERs are present in small amounts in membranes and play specialized regulatory functions in cellular metabolism, often depending on the chain length of their fatty acyl components. Particular attention was paid to CERs and their functions in the regulation of apoptosis as well as cell differentiation, transformation and proliferation [44].

**Fig. 6** displays circular charts that compare all types of samples taken for analysis. There are 13 masses characteristic of the CER group. When analyzing the cells’ glucose concentrations, it is visible that the change in culture conditions did not significantly affect the changes in the content of CERs. The only noticeable differences apply to NG and HG in the case of [C_32_H_61_NO_3_]^+^ - Cer 32:2 (*m/z* 508,72), for which in HG conditions, the relative content decreased by 2.6%. For other characteristic masses, the changes are less than 1%. The intensity values for the mentioned mass also differed in the case of large and small EVs. HG caused an increase in the content of this CER for both masses. In the case of large EVs, differences were also demonstrated for [C_41_H_77_NO_3_]^+^ - Cer 41:3, the content of which decreased with the increased glucose level, and for [C_42_H_81_NO_3_]^+^ Cer 42:2, for which the content increased. For small EVs, the largest differences were observed for [C_42_H_81_NO_3_]^+^ - Cer 42:2, for which, as in large EVs, the content increased significantly. For other characteristic ions, differences of up to 2% were observed.

**Fig. 6.**
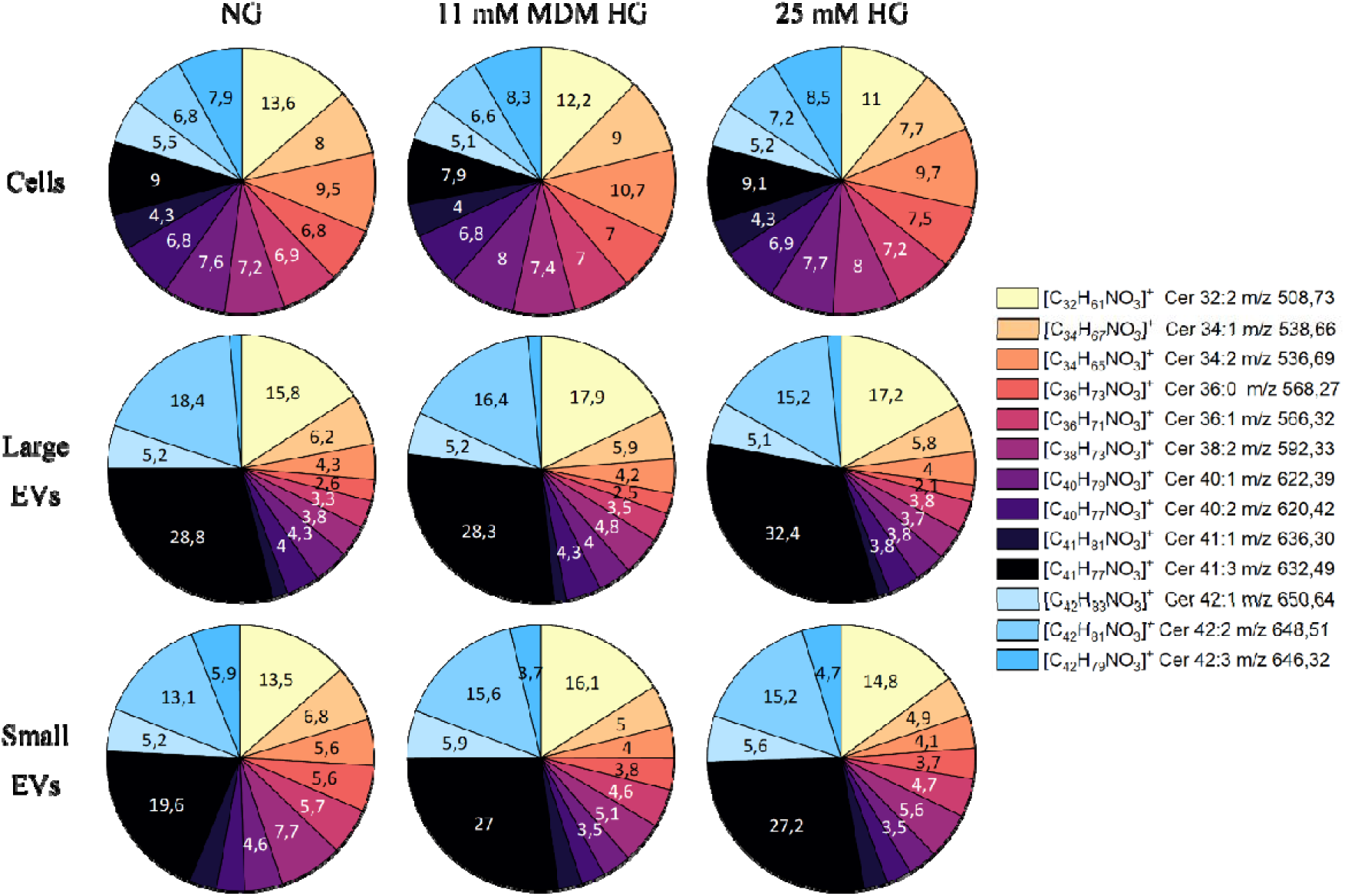
The percentages comparison calculated from the mean, normalized ion intensities of characteristic peaks in positive polarity for ceramides (CER) in the case of normoglycemia (NG), moderate hyperglycemia (11 mM MDM HG) and hyperglycemia (25 mM HG).

**Fig. 7** illustrates changes in the content of the CER group between cells, large and small EVs. The charts reveal that large and small EVs are significantly enriched in [C_34_H_67_NO_3_]^+^ - Cer 34:1, [C_36_H_73_NO_3_]^+^ - Cer 36:0 and [C_42_H_79_NO_3_]^+^ - Cer 42:3 compared to the cells from which they were isolated. The remaining masses characteristic of the CER group have minor changes of less than 4%. The same pattern was observed for the other two cultured conditions.

**Fig. 7.**
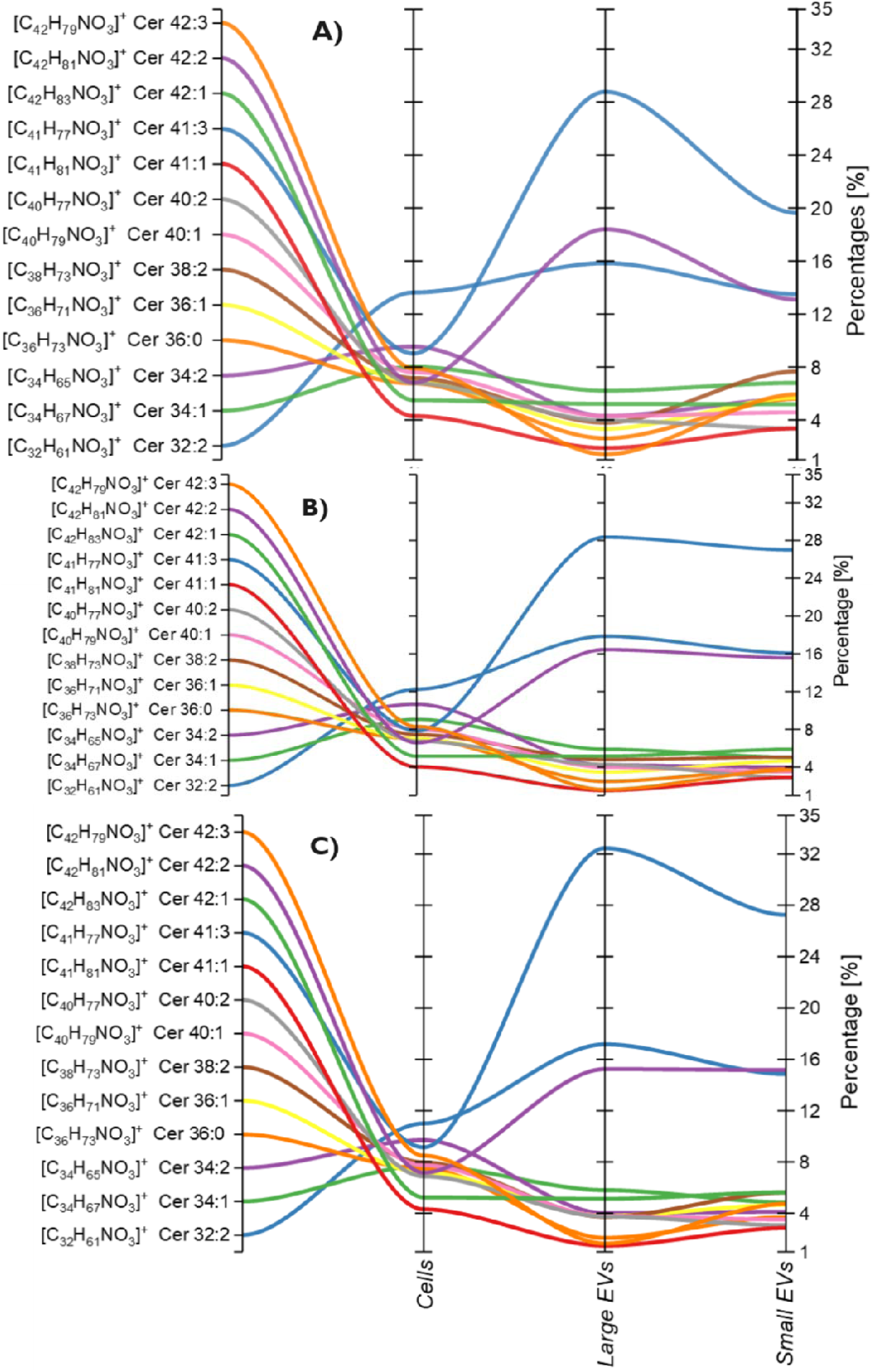
Percentage content changes for CER group in A) normoglycemic cells (5 mM D-glucose), B) moderately glycemic cells (11 mM glucose), and C) hyperglycemic cells (25 mM D-glucose).

##### a) Hexosylceramides (HexCER)

HexCER is a type of glycosphingolipid consisting of a ceramide backbone linked to a neutral sugar molecule. The different combination of its components results in a variety of monohexosylceramides, for example, 56 of them have been isolated from human plasma, which gives a greater variety in terms of CER, but a much smaller amount [45,46]. They are usually key precursors for the biosynthesis of more complex glycosphingolipids, including globosides and gangliosides, and are also structural components of cell membranes and lipid rafts [47].

In **Fig. 8**, the analysis performed did not reveal any changes in the β-cell culture caused by hyperglycemic conditions. However, both large and small EVs showed a slight increase in the content of [C_42_H_83_NO_8_]^+^ HexCer 36:0 and [C_42_H_81_NO_8_]^+^ HexCer 36:1, along with a simultaneous decrease in the content of [C_42_H_79_NO_8_]^+^ HexCer 36:2.

**Fig. 8.**
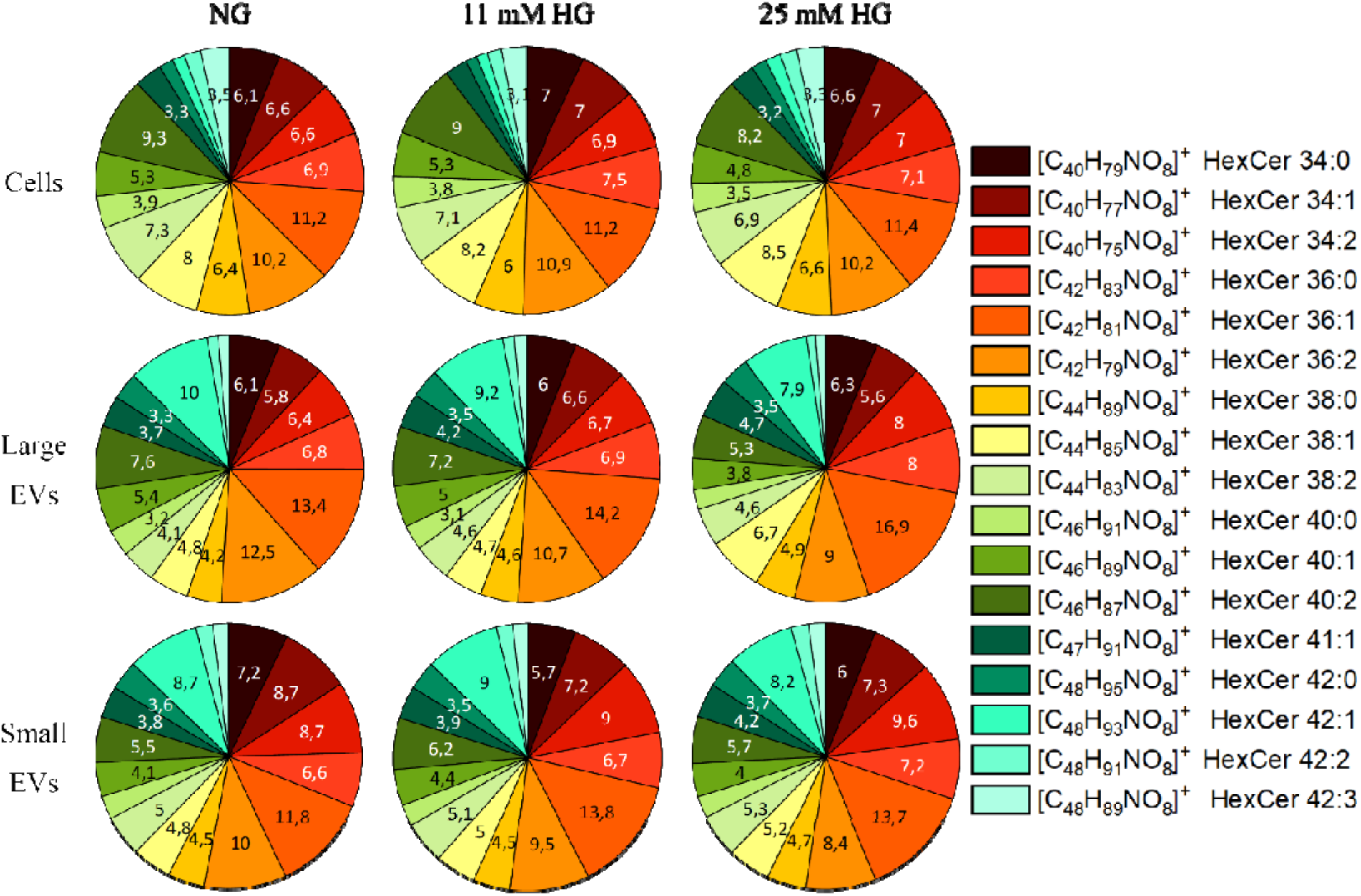
The percentages comparison calculated from the mean, normalized ion intensities of characteristic peaks in positive polarity for hexceramides (hexCER) in normoglycemia (NG), moderate hyperglycemia (11 mM MGM) and hyperglycemia (25 mM HG).

The analysis involved examining 17 peaks characteristic of hexCER with different acid lengths and different numbers of double bonds. The results revealed several differences recurrent for all three culture conditions, when comparing cells, large and small EVs (**Fig. 9**). A decrease in the content of: [C_44_H_89_NO_8_]^+^ HexCer 38:0 (*m/z* 760.62), [C_44_H_85_NO_8_]^+^ HexCer 38:1 (*m/z* 756.60), [C_44_H_83_NO_8_]^+^ HexCer 38:2 (*m/z* 754.54), [C_46_H_87_NO_8_]^+^ HexCer 40:2 (*m/z* 782.60) was observed in all EV samples, along with a simultaneous increase in the content of [C_48_H_93_NO_8_]^+^ HexCer 42:1 (*m/z* 812.62) and [C_42_H_81_NO_8_]^+^ HexCer 36:1 (*m/z* 728.51). The content of the remaining lipids from the hexCER group changed insignificantly by approximately 1% or remained at the same level.

**Fig. 9.**
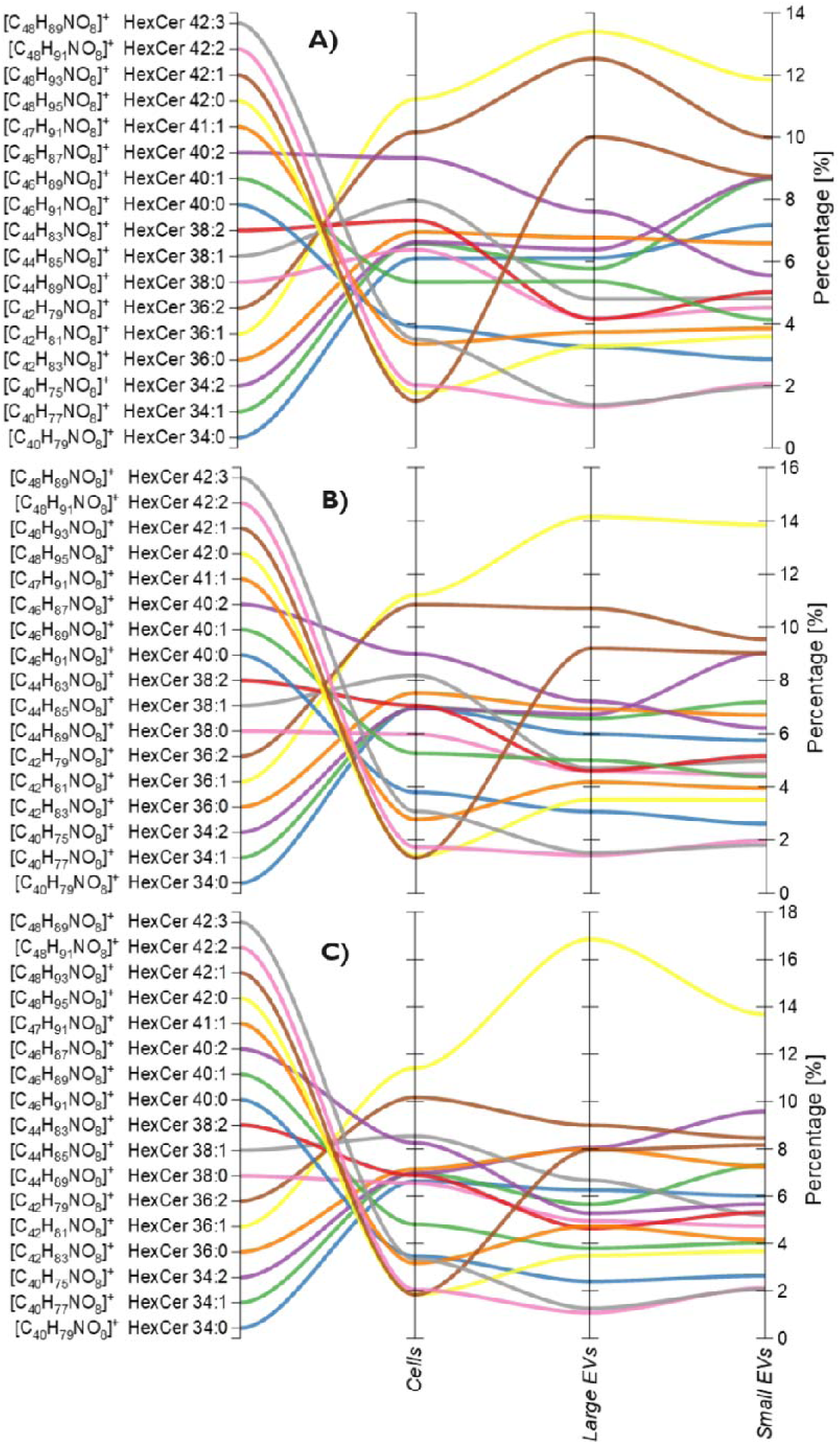
The percentage content changes for the hexCER group under three different conditions: A) normoglycemic cells (5 mM D-glucose), B) moderately glycemic cells (11 mM glucose), C) hyperglycemic cells (25 mM D-glucose).

##### b) Glycosphingolipids (GSLs)

GSLs are a type of lipid that consist of a hydrophilic carbohydrate chain and a hydrophobic ceramide. The carbohydrate chain acts as a ligand for carbohydrate-recognizing molecules, while the ceramide moiety forms an anchoring domain in the outer part of the cell membrane [48–50]. The region allows membrane proteins to function as receptors, channels and transporters. GSLs are critical in microdomain formation due to their ability to form systems through hydrogen bonds by donor and acceptor functional groups in ceramides, as well as hydrophobic interactions between their long-saturated acyl chains. They are also found in membrane rafts containing detergent-insoluble membrane components and signaling molecules [51,52]. Moreover, GSL is a group of compounds that differ in composition, quantity and distribution across species, tissues, and cells. In cells, this type of compounds most often occurs as an integral component of cell membranes, especially extracellular surface membranes.

GSLs are involved in a variety of physiological functions. Studies have shown that GSLs also take part in the processes that create inflammation, such as cancer and Alzheimer’s disease. Therefore, it is essential to discover additional information about this group of compounds [53–57].

**Fig. 10** shows the results of the analysis of 7 masses in the GSL group in relation to β-cells, large and small EVs Changes in this lipid group were observed in both culture conditions and sample type. Under HG conditions, there was a change in proportions for the cell, resulting in a significant increase in the content of [C_42_H_81_NO_3_Na]^+^ GSL molecular ion 18:0 with Na (*m/z* 750.49), [C_42_H_81_NO_3_Na]^+^ sulfatide 42:2 fragment (*m/z* 670, 47), [C_47_H_91_NO_9_Na]^+^ GSL molecular ion (*m/z* 836.80) with a decrease in the content for [C_2_H_9_NPO_4_]^+^ phosphosphingolipid fragment (*m/z* 142.07).

**Fig. 10.**
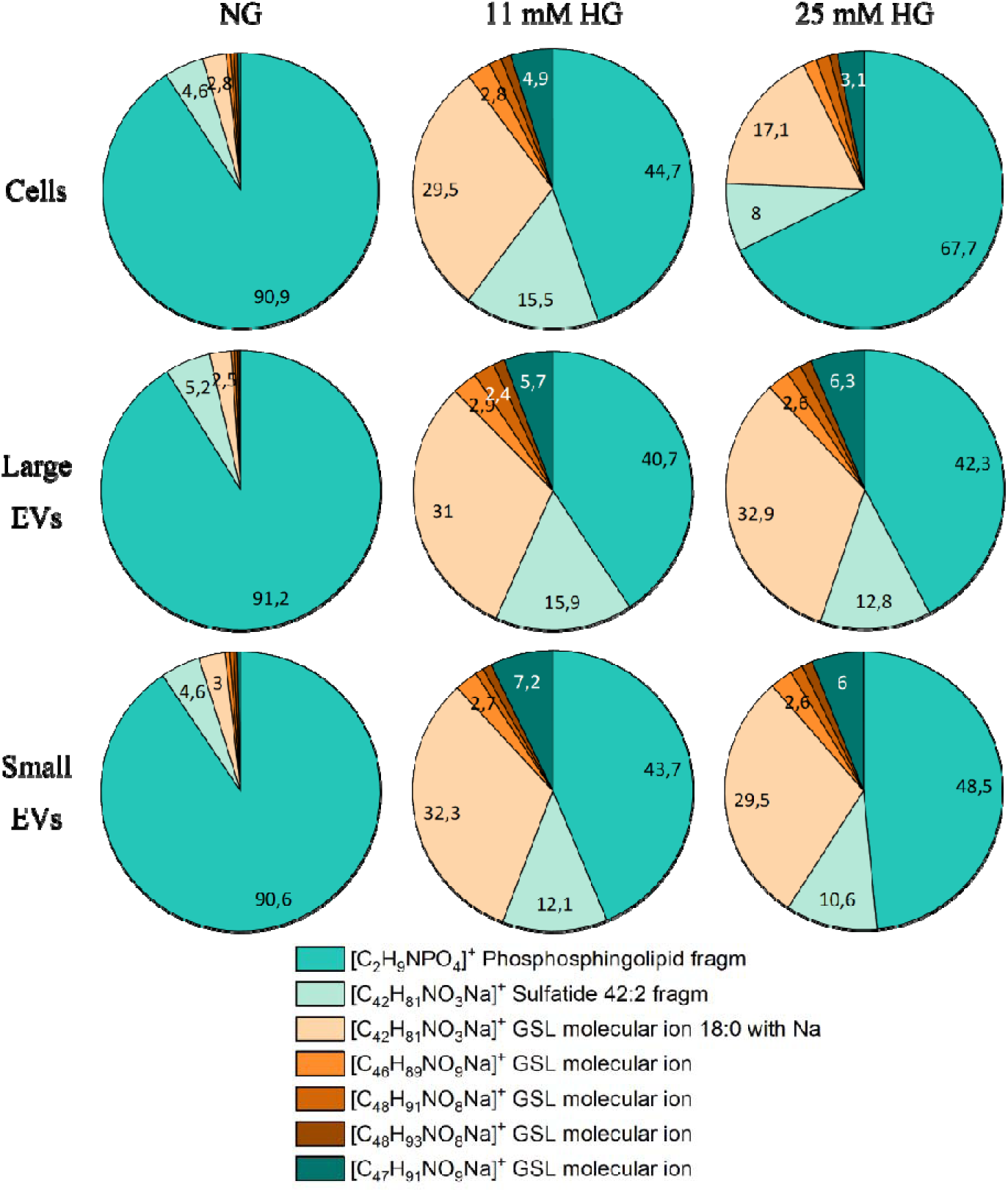
The percentages comparison calculated from the mean, normalized ion intensities of characteristic peaks in positive polarity for glycosphingolipids (GSLs) in normoglycemia (NG), moderate hyperglycemia (11 mM MGM) and hyperglycemia (25 mM HG).

Analyzing the results in terms of sample type (**Fig. 11**), there were no significant changes under NG conditions in the GLS ratio between cells, large and small EVs. However, for medium and high HG, an increase in the content of [C_47_H_91_NO_9_Na]^+^ GSL molecular ion (*m/z* 836.80) and [C_42_H_81_NO_3_Na]^+^ GSL molecular ion 18:0 with Na (*m/z* 750.49) in large and small ones can be observed.

**Fig. 11.**
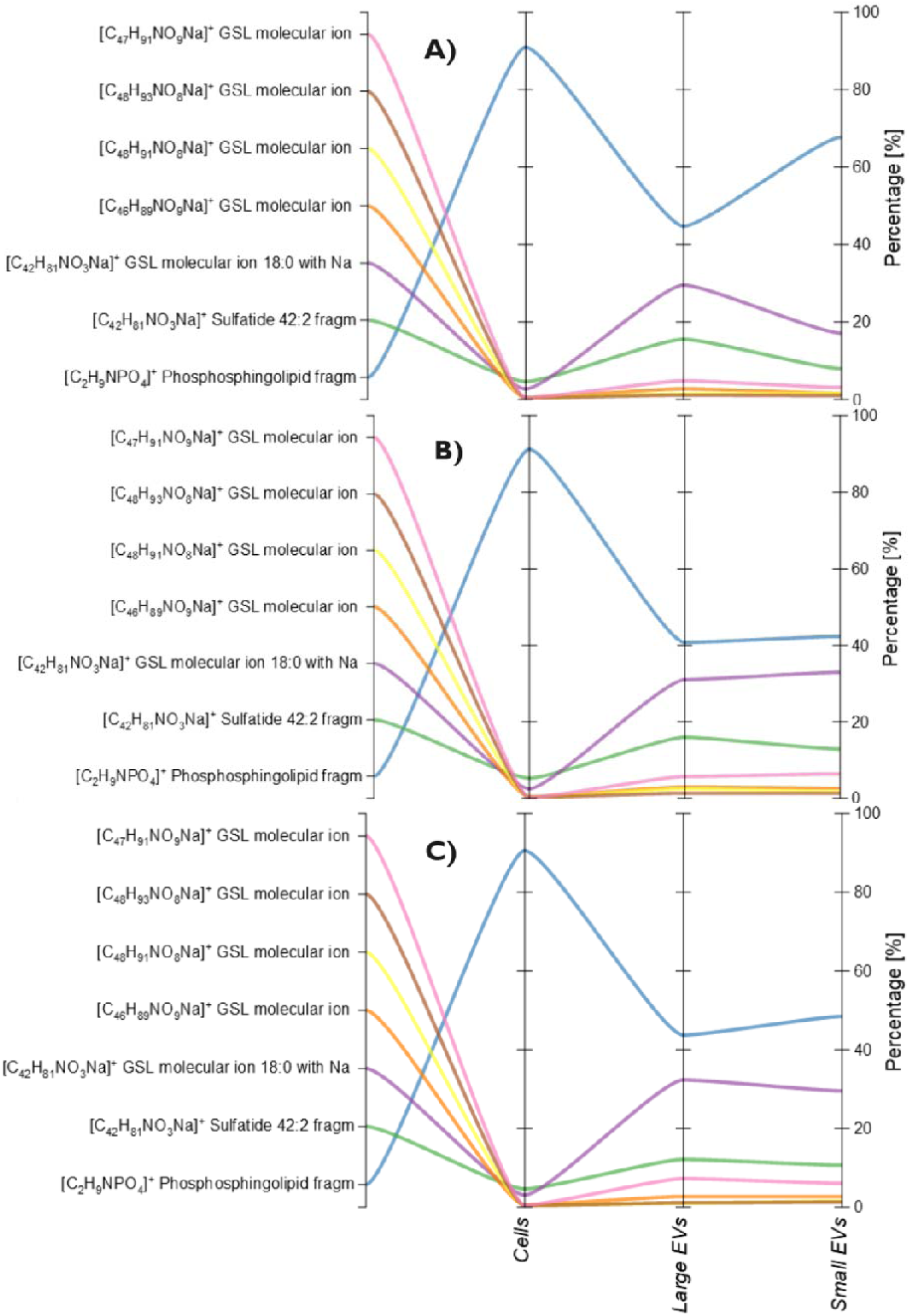
The percentage content changes in the glycosphingolipids group for A) normoglycemic cells (5 mM D-glucose), B) moderately glycemic cells (11 mM glucose), C) hyperglycemic cells (25 mM D-glucose).

## 4. Discussion

HG is a condition where there is too much sugar in the blood due to insufficient insulin to absorb and metabolize it. This condition mainly affects people with diabetes and can lead to serious long-term consequences. Over time, the elevated glucose levels disrupt the sugar metabolism and alter the physiology of the biological system. Pancreatic β-cells are one such biological system that secretes EVs, which are transporters of information. The information transfer happens at various levels, such as the change in the amount and presence of molecular compounds. An example of such compounds are SPs, which occur in cells as well as EVs. CER, HexCER and GSLs are lipids that are part of the complexes responsible for molecular processes inside the cell, and they can also serve as informative molecules reflecting changes in the presence and amount of pathological states. It is hypothesized that their presence might signal changes resulting from HG. The results indicate a significant decrease in the content of selected lipids from the mentioned classes due to the HG.

Previous research have shown that small EVs are enriched with specific lipids, i.e. glycosphingolipids, sphingomyelin, cholesterol and phosphatidylserine [23]. Among the proteins enriched in HG-induced small EVs are the participating proteins in active vesicular transport, such as IST1, ESCRT-III complex binding factor, SNARE proteins (VAMP1), as well as Ca^2+^-dependent phospholipid binding proteins (ANAXA11, CPNE8) or sterols (HPCAL1), HDL receptor proteins (SCRAB1) [58]. These proteins are actively involved in the lipid transport and their increased content in HG small EVs may be related to selective lipids enrichment. The comparative analysis carried out for this study confirmed that the small subpopulation of EVs is characterized by higher levels of mean, normalized intensity values in relation to the cells from which they originate. Moreover, the lipid profile depends on both the type of cell and its physiological state. Summing up, EVs from healthy cells may differ in composition from those cultured in conditions of disturbed physiology, e.g. HG. There are also differences in the composition of CER, HexCER, and GSL between departmental subpopulations of EVs from a specific cell line [59]. Ceramide-containing lipids play crucial roles in the human body and can be involved in neurodegenerative diseases such as Alzheimer’s and Parkinson’s. This study considers hyperglycemia and its impact on the level of ceramide-related lipids [60,61]. The issue is important because it is hypothesized that the action of ceramide on the content and biogenesis of EVs in cells may have a similar effect on ceramides located on EVs. Therefore, EVs enriched with ceramides may initiate apoptosis in astrocytes, and a similar mechanism may occur in HG [62].

Further studies extend the known role of ceramides, hexceramides and glycosphingolipids in the development of pathological processes using a novel approach. To achieve this, new methods of identification and analysis need to be employed.

## 5. Conclusion

The study confirmed the presence of both large and small EVs in isolated samples through various analyses, including cryo-TEM and TEM, and spectral flow cytometry. The detection of specific markers like CD9, CD63 and annexin-V helped in the confirmation. As demonstrated by Mathilde Mathieu et al., both tetraspanins CD9 and CD63 can be released in small ectosomes formed at the plasma membrane, a finding that our results corroborate [63]. Similar observations were reported in studies performed on the immortalized endothelial cells [64].

Utilizing ToF-SIMS mass spectrometry, we performed a comparative analysis of 38 lipids belonging to the sphingolipid group. The percentage of content of mean and normalized intensity values were compared for the experimental groups, including *β*-cells, large and small extracellular vesicles from normoglycemia, medium glycemia and hyperglycemia. The examination of EVs’ content and distribution with ToF-SIMS can provide insight into the physiopathology associated with diabetes and contribute to the discovery of new biomarkers for early diagnosis and risk. This work focuses on lipids and their informative function in communication *via* extracellular vesicles, while it has been shown that transported cargo EVs, such as mRNA and proteins play such a role [65,66]. To the best of our knowledge, this is the first study to show TOF-SIMS results, including a comparative analysis of ceramides, hexolysceramides and glycosphingolipids in extracellular vesicles derived from *β*-cells cultured in hyperglycemia.

## Data Availability

The data underlying this article are available at RODBUK Jagiellonian University in Krakow at https://dx.doi.org/10.57903/UJ/VJKZDE.

## Author contributions

M.S. – data curation, formal analysis, investigation, methodology, validation, visualisation, writing – original draft; M. D.-K. – investigation, methodology, validation, visualisation, writing original draft; E.Ł.S. conceptualization, formal analysis, funding acquisition, methodology, project administration, resources, supervision, writing – review & editing

## Acknowledgements

1. This publication has been funded from the SciMat and qLife Priority Research Area budget under the Strategic Programme Excellence Initiative at the Jagiellonian University.
2. This work was supported by the National Science Center (NCN), grant OPUS 17 to Prof. E. Stępień (No. 2019/33/B/NZ3/01004).
3. This publication was developed under the provision of the Polish Ministry and Higher Education project Support for research and development with the use of research infrastructure of the National Synchrotron Radiation Centre SOLARIS” under contract nr 1/SOL/2021/2. We acknowledge the SOLARIS Centre for the access to the cryogenic electron microscope, where the measurements were performed.
4. The authors would like to thank Dr. Olga Barczyk-Woźnicka from the Institute of Zoology and Biomedical Research of the Jagiellonian University for providing TEM images. The authors would like to thank Dr Michał Rawski from the SOLARIS National Synchrotron Radiation Center for providing cryo-TEM images.The authors would like to thank Olena Bohomolova for her support during EVs isolation.

## References

[1] W. Zheng, J. Kollmeyer, H. Symolon, A. Momin, E. Munter, E. Wang, S. Kelly, J.C. Allegood, Y. Liu, Q. Peng, H. Ramaraju, M.C. Sullards, M. Cabot, A.H. Merrill, Ceramides and other bioactive sphingolipid backbones in health and disease: Lipidomic analysis, metabolism and roles in membrane structure, dynamics, signaling and autophagy, Biochimica et Biophysica Acta (BBA) - Biomembranes 1758 (2006) 1864–1884. 10.1016/j.bbamem.2006.08.009.

[2] A.H.J. Merrill, T.H. Stokes, A. Momin, H. Park, B.J. Portz, S. Kelly, E. Wang, M.C. Sullards, M.D. Wang, Sphingolipidomics: a valuable tool for understanding the roles of sphingolipids in biology and disease., Journal of Lipid Research 50 Suppl (2009) S97–102. 10.1194/jlr.R800073-JLR200.

[3] M.P. Stępień EŁ, Durak-Kozica M, Extracellular vesicles in vascular pathophysiology: beyond their molecular content, Pol Arch Intern Med 133 (2023) 16483. doi:10.20452/pamw.16483.

[4] M.K. Cavaghan, D.A. Ehrmann, K.S. Polonsky, Interactions between insulin resistance and insulin secretion in the development of glucose intolerance., The Journal of Clinical Investigation 106 (2000) 329–333. 10.1172/JCI10761.

[5] M.E. Cerf, Beta cell dysfunction and insulin resistance., Frontiers in Endocrinology 4 (2013) 37. 10.3389/fendo.2013.00037.

[6] J.L. Leahy, Pathogenesis of Type 2 Diabetes Mellitus, Archives of Medical Research 36 (2005) 197–209. 10.1016/j.arcmed.2005.01.003.

[7] M.T. Malecki, Type 2 diabetes mellitus and its complications: from the molecular biology to the clinical practice., The Review of Diabetic Studieslll: RDS 1 (2004) 5–8. 10.1900/RDS.2004.1.5.

[8] T. Tomita, Apoptosis in pancreatic β-islet cells in Type 2 diabetes., Bosnian Journal of Basic Medical Sciences 16 (2016) 162–179. 10.17305/bjbms.2016.919.

[9] Z. Fu, E.R. Gilbert, D. Liu, Regulation of insulin synthesis and secretion and pancreatic Beta-cell dysfunction in diabetes, Curr Diabetes Rev 9 (2013) 25–53.

[10] S.B. Russo, J.S. Ross, L.A. Cowart, Sphingolipids in obesity, type 2 diabetes, and metabolic disease., Handb Exp Pharmacol (2013) 373–401. 10.1007/978-3-7091-1511-4_19.

[11] N. Tofte, T. Suvitaival, L. Ahonen, S.A. Winther, S. Theilade, M. Frimodt-Møller, T.S. Ahluwalia, P. Rossing, Lipidomic analysis reveals sphingomyelin and phosphatidylcholine species associated with renal impairment and all-cause mortality in type 1 diabetes, Sci Rep 9 (2019) 16398. 10.1038/s41598-019-52916-w.

[12] M.E. Marzec, D. Wojtysiak, K. Połtowicz, J. Nowak, R. Pedrys, Study of cholesterol and vitamin E levels in broiler meat from different feeding regimens by TOF-SIMS, Biointerphases 11 (2016). 10.1116/1.4943619.

[13] D.C. Fernández-Remolar, D. Gómez-Ortiz, P. Malmberg, T. Huang, Y. Shen, A. Anglés, R. Amils, Preservation of Underground Microbial Diversity in Ancient Subsurface Deposits (>6 Ma) of the Rio Tinto Basement, Microorganisms 9 (2021). 10.3390/microorganisms9081592.

[14] T.B. Angerer, D. Velickovic, C.D. Nicora, J.E. Kyle, D.J. Graham, C. Anderton, L.J. Gamble, Exploiting the Semidestructive Nature of Gas Cluster Ion Beam Time-of-Flight Secondary Ion Mass Spectrometry Imaging for Simultaneous Localization and Confident Lipid Annotations, Anal Chem 91 (2019) 15073–15080. 10.1021/acs.analchem.9b03763.

[15] D.-K. Kim, J. Lee, S.R. Kim, D.-S. Choi, Y.J. Yoon, J.H. Kim, G. Go, D. Nhung, K. Hong, S.C. Jang, S.-H. Kim, K.-S. Park, O.Y. Kim, H.T. Park, J.H. Seo, E. Aikawa, M. Baj-Krzyworzeka, B.W.M. Van Balkom, M. Belting, L. Blanc, V. Bond, A. Bongiovanni, F.E. Borràs, L. Buée, E.I. Buzás, L. Cheng, A. Clayton, E. Cocucci, C.S. Dela Cruz, D.M. Desiderio, D. Di Vizio, K. Ekström, J.M. Falcon-Perez, C. Gardiner, B. Giebel, D.W. Greening, J. Christina Gross, D. Gupta, A. Hendrix, A.F. Hill, M.M. Hill, E. Nolte-’T Hoen, D.W. Hwang, J. Inal, M.V. Jagannadham, M. Jayachandran, Y.-K. Jee, M. Jørgensen, K.P. Kim, Y.-K. Kim, T. Kislinger, C. Lässer, D.S. Lee, H. Lee, J. Van Leeuwen, T. Lener, M.-L. Liu, J. Lötvall, A. Marcilla, S. Mathivanan, A. Möller, J. Morhayim, F. Mullier, I. Nazarenko, R. Nieuwland, D.N. Nunes, K. Pang, J. Park, T. Patel, G. Pocsfalvi, H. Del Portillo, U. Putz, M.I. Ramirez, M.L. Rodrigues, T.-Y. Roh, F. Royo, S. Sahoo, R. Schiffelers, S. Sharma, P. Siljander, R.J. Simpson, C. Soekmadji, P. Stahl, A. Stensballe, E. Stepień, H. Tahara, A. Trummer, H. Valadi, L.J. Vella, S.N. Wai, K. Witwer, M. Yánez-Mó, H. Youn, R. Zeidler, Y.S. Gho, EVpedia: A community web portal for extracellular vesicles research, Bioinformatics 31 (2015). 10.1093/bioinformatics/btu741.

[16] M. Durak-Kozica, A. Wróbel, M. Platt, E.Ł. Stępień, Comparison of qNANO results from the isolation of extracellular microvesicles with the theoretical model, Bio-Algorithms and Med-Systems 18 (2022) 171–179. doi:10.2478/bioal-2022-0088.

[17] K.W. Witwer, C. Théry, Extracellular vesicles or exosomes? On primacy, precision, and popularity influencing a choice of nomenclature, J Extracell Vesicles 8 (2019) 1648167. 10.1080/20013078.2019.1648167.

[18] E.Ł. Stlllpien, M. Durak-Kozica, A. Kamińska, M. Targosz-Korecka, M. Libera, G. Tylko, A. Opalińska, M. Kapusta, B. Solnica, A. Georgescu, M.C. Costa, A. Czyzewska-Buczyńska, W. Witkiewicz, M.T. Malecki, F.J. Enguita, Circulating ectosomes: Determination of angiogenic microRNAs in type 2 diabetes, Theranostics 8 (2018). 10.7150/thno.23334.

[19] M. Yáñez-Mó, P.R.-M. Siljander, Z. Andreu, A.B. Zavec, F.E. Borràs, E.I. Buzas, K. Buzas, E. Casal, F. Cappello, J. Carvalho, E. Colás, A. Cordeiro-da Silva, S. Fais, J.M. Falcon-Perez, I.M. Ghobrial, B. Giebel, M. Gimona, M. Graner, I. Gursel, M. Gursel, N.H.H. Heegaard, A. Hendrix, P. Kierulf, K. Kokubun, M. Kosanovic, V. Kralj-Iglic, E.-M. Krämer-Albers, S. Laitinen, C. Lässer, T. Lener, E. Ligeti, A. Linē, G. Lipps, A. Llorente, J. Lötvall, M. Manček-Keber, A. Marcilla, M. Mittelbrunn, I. Nazarenko, E.N.M. Nolte-’t Hoen, T.A. Nyman, L. O’Driscoll, M. Olivan, C. Oliveira, É. Pállinger, H.A. Del Portillo, J. Reventós, M. Rigau, E. Rohde, M. Sammar, F. Sánchez-Madrid, N. Santarém, K. Schallmoser, M.S. Ostenfeld, W. Stoorvogel, R. Stukelj, S.G. Van der Grein, M.H. Vasconcelos, M.H.M. Wauben, O. De Wever, Biological properties of extracellular vesicles and their physiological functions, J Extracell Vesicles 4 (2015) 27066. 10.3402/jev.v4.27066.

[20] M.A. Di Bella, Overview and Update on Extracellular Vesicles: Considerations on Exosomes and Their Application in Modern Medicine., Biology (Basel) 11 (2022). 10.3390/biology11060804.

[21] E.Ł. Stępień, A. Kamińska, M. Surman, D. Karbowska, A. Wróbel, M. Przybyło, Fourier-Transform InfraRed (FT-IR) spectroscopy to show alterations in molecular composition of EV subpopulations from melanoma cell lines in different malignancy, Biochemistry and Biophysics Reports 25 (2021) 100888. 10.1016/j.bbrep.2020.100888.

[22] C. Rząca, U. Jankowska, E.Ł. Stępień, Proteomic profiling of exosomes derived from pancreatic beta-cells cultured under hyperglycemia, Bio-Algorithms and Med-Systems 18 (2022) 151– 157. doi:10.2478/bioal-2022-0085.

[23] A. Llorente, T. Skotland, T. Sylvänne, D. Kauhanen, T. Róg, A. Orłowski, I. Vattulainen, K. Ekroos, K. Sandvig, Molecular lipidomics of exosomes released by PC-3 prostate cancer cells, Biochimica et Biophysica Acta (BBA) - Molecular and Cell Biology of Lipids 1831 (2013) 1302– 1309. 10.1016/j.bbalip.2013.04.011.

[24] K. Carayon, K. Chaoui, E. Ronzier, I. Lazar, J. Bertrand-Michel, V. Roques, S. Balor, F. Terce, A. Lopez, L. Salomé, E. Joly, Proteolipidic Composition of Exosomes Changes during Reticulocyte Maturation*, Journal of Biological Chemistry 286 (2011) 34426–34439. 10.1074/jbc.M111.257444.

25. [25] L. Musante, D. Tataruch, D. Gu, A. Benito-Martin, G. Calzaferri, S. Aherne, H. Holthofer, A Simplified Method to Recover Urinary Vesicles for Clinical Applications and Sample Banking, Scientific Reports 4 (2014) 7532. 10.1038/srep07532.

[26] A. Kamińska, M. Roman, A. Wróbel, A. Gala-Błądzińska, M.T. Małecki, C. Paluszkiewicz, E.Ł. Stępień, Raman spectroscopy of urinary extracellular vesicles to stratify patients with chronic kidney disease in type 2 diabetes, Nanomedicine: Nanotechnology, Biology and Medicine 39 (2022) 102468. 10.1016/j.nano.2021.102468.

[27] M.E. Marzec, C. Rząca, P. Moskal, E. Stępień, Study of the influence of hyperglycemia on the abundance of amino acids, fatty acids, and selected lipids in extracellular vesicles using TOF-SIMS, Biochem Biophys Res Commun 622 (2022) 30–36. 10.1016/j.bbrc.2022.07.020.

[28] M.E. Marzec, D. Wojtysiak, K. Połtowicz, J. Nowak, R. Pedrys, Study of cholesterol and vitamin E levels in broiler meat from different feeding regimens by TOF-SIMS, Biointerphases 11 (2016) 02A326. 10.1116/1.4943619.

[29] A. Kamińska, M.E. Marzec, E.Ł. Stępień, Design and Optimization of a Biosensor Surface Functionalization to Effectively Capture Urinary Extracellular Vesicles., Molecules 26 (2021). 10.3390/molecules26164764.

[30] D.J. Graham, D.G. Castner, Image and Spectral Processing for ToF-SIMS Analysis of Biological Materials., Mass Spectrom (Tokyo) 2 (2013) S0014. 10.5702/massspectrometry.S0014.

[31] B.J. Tyler, G. Rayal, D.G. Castner, Multivariate analysis strategies for processing ToF-SIMS images of biomaterials., Biomaterials 28 (2007) 2412–2423. 10.1016/j.biomaterials.2007.02.002.

[32] M. Skalska, M. Durak-Kozica, Experimental and analytical procedures for the ToF-SIMS measurement data of membranous structures., Bio-Algorithms and Med-Systems 19 (2023) 64–68. 10.5604/01.3001.0054.1935.

[33] S. Hallal, Á. Tűzesi, G.E. Grau, M.E. Buckland, K.L. Alexander, Understanding the extracellular vesicle surface for clinical molecular biology., Journal of Extracellular Vesicles 11 (2022) e12260. 10.1002/jev2.12260.

[34] Y. Zhang, Y. Liu, H. Liu, W.H. Tang, Exosomes: biogenesis, biologic function and clinical potential, Cell & Bioscience 9 (2019) 19. 10.1186/s13578-019-0282-2.

[35] S. Keerthikumar, D. Chisanga, D. Ariyaratne, H. Al Saffar, S. Anand, K. Zhao, M. Samuel, M. Pathan, M. Jois, N. Chilamkurti, L. Gangoda, S. Mathivanan, ExoCarta: A Web-Based Compendium of Exosomal Cargo., Journal of Molecular Biology 428 (2016) 688–692. 10.1016/j.jmb.2015.09.019.

[36] B.M. Castro, M. Prieto, L.C. Silva, Ceramide: A simple sphingolipid with unique biophysical properties, Progress in Lipid Research 54 (2014) 53–67. 10.1016/j.plipres.2014.01.004.

[37] T. Skotland, N.P. Hessvik, K. Sandvig, A. Llorente, Exosomal lipid composition and the role of ether lipids and phosphoinositides in exosome biology, J Lipid Res 60 (2019) 9–18. 10.1194/jlr.R084343.

[38] L. Salimi, A. Akbari, N. Jabbari, B. Mojarad, A. Vahhabi, S. Szafert, S.A. Kalashani, H. Soraya, M. Nawaz, J. Rezaie, Synergies in exosomes and autophagy pathways for cellular homeostasis and metastasis of tumor cells, Cell Biosci 10 (2020) 64. 10.1186/s13578-020-00426-y.

[39] M. Record, S. Silvente-Poirot, M. Poirot, M.J.O. Wakelam, Extracellular vesicles: Lipids as key components of their biogenesis and functions, J Lipid Res 59 (2018) 1316–1324. 10.1194/jlr.E086173.

[40] T. Matsui, F. Osaki, S. Hiragi, Y. Sakamaki, M. Fukuda, ALIX and ceramide differentially control polarized small extracellular vesicle release from epithelial cells, EMBO Rep 22 (2021) e51475. 10.15252/embr.202051475.

[41] G.O. Skryabin, A. V Komelkov, E.E. Savelyeva, E.M. Tchevkina, Lipid Rafts in Exosome Biogenesis, Biochemistry (Moscow) 85 (2020) 177–191. 10.1134/S0006297920020054.

[42] E. Bieberich, Sphingolipids and lipid rafts: Novel concepts and methods of analysis., Chemistry and Physics of Lipids 216 (2018) 114–131. 10.1016/j.chemphyslip.2018.08.003.

[43] P. Sigmund, Theory of Sputtering. I. Sputtering Yield of Amorphous and Polycrystalline Targets, Physical Review 184 (1969) 383–416.

[44] C.C. Geilen, T. Wieder, C.E. Orfanos, Ceramide signalling: regulatory role in cell proliferation, differentiation and apoptosis in human epidermis., Arch Dermatol Res 289 (1997) 559–566. 10.1007/s004030050240.

[45] O. Quehenberger, A.M. Armando, A.H. Brown, S.B. Milne, D.S. Myers, A.H. Merrill, S. Bandyopadhyay, K.N. Jones, S. Kelly, R.L. Shaner, C.M. Sullards, E. Wang, R.C. Murphy, R.M. Barkley, T.J. Leiker, C.R.H. Raetz, Z. Guan, G.M. Laird, D.A. Six, D.W. Russell, J.G. McDonald, S. Subramaniam, E. Fahy, E.A. Dennis, Lipidomics reveals a remarkable diversity of lipids in human plasma1[S], J Lipid Res 51 (2010) 3299–3305. 10.1194/jlr.M009449.

[46] B.W.S. Lam, T.Y.A. Yam, C.P. Chen, M.K.P. Lai, W.-Y. Ong, D.R. Herr, The noncanonical chronicles: Emerging roles of sphingolipid structural variants, Cell Signal 79 (2021) 109890. 10.1016/j.cellsig.2020.109890.

[47] U. Loizides-Mangold, F.P.A. David, V.J. Nesatyy, T. Kinoshita, H. Riezman, Glycosylphosphatidylinositol anchors regulate glycosphingolipid levels., J Lipid Res 53 (2012) 1522–1534. 10.1194/jlr.M025692.

[48] S. Hakomori, Carbohydrate-to-carbohydrate interaction in basic cell biology: a brief overview., Archives of Biochemistry and Biophysics 426 (2004) 173–181. 10.1016/j.abb.2004.02.032.

[49] A. Regina Todeschini, S. Hakomori, Functional role of glycosphingolipids and gangliosides in control of cell adhesion, motility, and growth, through glycosynaptic microdomains., Biochimica et Biophysica Acta 1780 (2008) 421–433. 10.1016/j.bbagen.2007.10.008.

[50] S. Sonnino, L. Mauri, V. Chigorno, A. Prinetti, Gangliosides as components of lipid membrane domains, Glycobiology 17 (2007) 1R–13R. 10.1093/glycob/cwl052.

[51] G. Gupta, A. Surolia, Glycosphingolipids in microdomain formation and their spatial organization., FEBS Lett 584 (2010) 1634–1641. 10.1016/j.febslet.2009.11.070.

[52] N.M. Hooper, Detergent-insoluble glycosphingolipid/cholesterol-rich membrane domains, lipid rafts and caveolae (review)., Mol Membr Biol 16 (1999) 145–156. 10.1080/096876899294607.

[53] C.A. Lingwood, Glycosphingolipid functions., Cold Spring Harb Perspect Biol 3 (2011). 10.1101/cshperspect.a004788.

[54] G. D’Angelo, S. Capasso, L. Sticco, D. Russo, Glycosphingolipids: synthesis and functions., FEBS J 280 (2013) 6338–6353. 10.1111/febs.12559.

[55] Z. Vukelić, S. Kalanj-Bognar, M. Froesch, L. Bîndila, B. Radić, M. Allen, J. Peter-Katalinić, A.D. Zamfir, Human gliosarcoma-associated ganglioside composition is complex and distinctive as evidenced by high-performance mass spectrometric determination and structural characterization., Glycobiology 17 (2007) 504–515. 10.1093/glycob/cwm012.

[56] D.D. Park, G. Xu, M. Wong, C. Phoomak, M. Liu, N.E. Haigh, S. Wongkham, P. Yang, E. Maverakis, C.B. Lebrilla, Membrane glycomics reveal heterogeneity and quantitative distribution of cell surface sialylation., Chem Sci 9 (2018) 6271–6285. 10.1039/c8sc01875h.

[57] G. van Echten-Deckert, J. Walter, Sphingolipids: critical players in Alzheimer’s disease., Prog Lipid Res 51 (2012) 378–393. 10.1016/j.plipres.2012.07.001.

[58] C. Rząca, U. Jankowska, E.Ł. Stępień, Proteomic profiling of exosomes derived from pancreatic beta-cells cultured under hyperglycemia, Bio-Algorithms and Med-Systems 18 (2022) 151– 157. 10.2478/bioal-2022-0085.

[59] J.F. Brouwers, M. Aalberts, J.W.A. Jansen, G. van Niel, M.H. Wauben, T.A.E. Stout, J.B. Helms, W. Stoorvogel, Distinct lipid compositions of two types of human prostasomes, Proteomics 13 (2013) 1660–1666. 10.1002/pmic.201200348.

[60] N. Bengoa-Vergniory, R.F. Roberts, R. Wade-Martins, J. Alegre-Abarrategui, Alpha-synuclein oligomers: a new hope, Acta Neuropathol 134 (2017) 819–838. 10.1007/s00401-017-1755-1.

[61] M.B. Dinkins, G. Wang, E. Bieberich, Sphingolipid-Enriched Extracellular Vesicles and Alzheimer’s Disease: A Decade of Research, Journal of Alzheimer’s Disease 60 (2017) 757–768. 10.3233/JAD-160567.

[62] G. Wang, M. Dinkins, Q. He, G. Zhu, C. Poirier, A. Campbell, M. Mayer-Proschel, E. Bieberich, Astrocytes Secrete Exosomes Enriched with Proapoptotic Ceramide and Prostate Apoptosis Response 4 (PAR-4): POTENTIAL MECHANISM OF APOPTOSIS INDUCTION IN ALZHEIMER DISEASE (AD)*, Journal of Biological Chemistry 287 (2012) 21384–21395. 10.1074/jbc.M112.340513.

[63] M. Mathieu, N. Névo, M. Jouve, J.I. Valenzuela, M. Maurin, F.J. Verweij, R. Palmulli, D. Lankar, F. Dingli, D. Loew, E. Rubinstein, G. Boncompain, F. Perez, C. Théry, Specificities of exosome versus small ectosome secretion revealed by live intracellular tracking of CD63 and CD9., Nat Commun 12 (2021) 4389. 10.1038/s41467-021-24384-2.

[64] M. Durak-Kozica, A. Wróbel, M. Platt, E.Ł. Stępień, Comparison of qNANO results from the isolation of extracellular microvesicles with the theoretical model, Bio-Algorithms and Med-Systems 18 (2022) 171–179. 10.2478/bioal-2022-0088.

[65] G.E. Grieco, D. Fignani, C. Formichi, L. Nigi, G. Licata, C. Maccora, N. Brusco, G. Sebastiani, F. Dotta, Extracellular Vesicles in Immune System Regulation and Type 1 Diabetes: Cell-to-Cell Communication Mediators, Disease Biomarkers, and Promising Therapeutic Tools, Frontiers in Immunology 12 (2021).

[66] K.R. Giri, L. de Beaurepaire, D. Jegou, M. Lavy, M. Mosser, A. Dupont, R. Fleurisson, L. Dubreil, M. Collot, P. Van Endert, J.-M. Bach, G. Mignot, S. Bosch, Molecular and Functional Diversity of Distinct Subpopulations of the Stressed Insulin-Secreting Cell’s Vesiculome, Frontiers in Immunology 11 (2020).

